# PBAE-PEG based lipid nanoparticles for lung cell-specific gene delivery

**DOI:** 10.1101/2024.08.22.608643

**Authors:** Bingxin Liu, Yamato Sajiki, Anusha Sridharan, William D. Stuart, Xuemei Cui, Matthew Siefert, Koichi Araki, Assem G. Ziady, Donglu Shi, Jeffery A. Whitsett, Yutaka Maeda

## Abstract

Delivery of modified mRNA encapsulated in lipid nanoparticles, exemplified by their successful use in COVID-19 vaccination, provides a framework for treating various genetic and acquired disorders. Herein, we developed PEGylated(PBAE-PEG) and non-PEGylated(PBAE) PBAE with lipids 4A3-SC8/DOPE/cholesterol/DOTAP to form lipid nanoparticles (LNPs) for mRNA delivery into different types of pulmonary cells in vivo. PBAE-PEG/LNP were highly active in transfecting HEK293T cells and air-liquid interfaced H441 cells in vitro. PBAE-PEG/LNP were used to express Cre-recombinase after administration to mice by intravenous injection, resulting in high transfection levels in pulmonary vascular endothelial cells. Intratracheal injection of both PBAE-PEG/LNP and PBAE/LNPs resulted in efficient and selective transfection of lung epithelial cells, identified by the expression of stabilized Cre-recombinase mRNA in club cells and alveolar type 2 cells. PBAE-PEG/LNP were most effective in transfecting alveolar type 2 cells after intratracheal injection, while PBAE/LNPs administered intratracheally were more effective in transfecting secretory airway cells. Cre-mediated recombination was specific to lung epithelial cells after intratracheal administration. Likewise, intravenous administration resulted in selective transfection of endothelial cells but not other pulmonary cell types, indicating their failure to cross the pulmonary endothelial-to-epithelial barrier. Moreover, 5-methoxyuridine modified mRNA was more efficient than unmodified mRNA *in vivo* but not *in vitro*.

## Introduction

Genetic and acquired disorders affecting lung function contribute to the global burden of lung disease. Mutations in genes critical for normal functions of pulmonary epithelial, endothelial, and mesenchymal cells cause a number of debilitating conditions, including cystic fibrosis, surfactant protein disorders, ciliary dyskinesia, and pulmonary phospholipid clearance disorders. Mutations in a number of genes, including SFTPC[1], SFTPB[2], NKX2-1[3] ABCA3[4], [5], CFTR[6], TBX4[7], and FOXF1[8], are essential for lung formation or function[9]. Mutations or dysregulation of these genes cause life-threatening diffuse pulmonary disorders in infants and children.

Over several decades, lentiviruses and adeno-associated viruses (AAV) have been actively studied for in vivo gene delivery to the lung [10][11][12] and have been used for transporting DNA[13], RNA[14], siRNA[15], and CRISPR-associated guide RNAs (gRNA)[12][16]. Despite their extensive study, immune responses that limit repeated administration challenge the application of viral vectors. While AAV vectors are being developed for delivery to pulmonary cells, clinical use of AAV for delivery to the human lung has not been achieved due to pre-existing immunity and the generation of immune responses. Safety concerns related to their integration into the host genome, risking insertional mutagenesis, have been raised.[15][17][18] These limitations underscore the need for developing high-efficiency, nonviral delivery platforms for the treatment of lung disease. A diversity of nonviral systems gene delivery, including nanoparticle-based methods, are being developed to mitigate immune and inflammatory responses to enable safer, repeatable treatments for various genetic disorders, including pulmonary diseases.

Recent advances in applying lung-targeted drug delivery with organ-targeting (SORT) lipids[19] and Poly(β-amino esters) (PBAE) have been summarized [20][21]. PBAE and SORT lipids exhibit low toxicity, biocompatibility, and biodegradability compared with PEI-based and PLL-based nanoparticles[22][23][24][25][26]; however, the combination of PBAE polymers with PEGylation (PBAE-PEG) and biocompatible lipids to create PBAE-LNP has not been extensively explored. In the present study, we synthesized two distinct types of PBAEs, PBAE-PEG and PBAE. We combined them with a lipid mixture comprising DOPE/DOTAP/4A3-SC8/Cholesterol to produce two unique nanoparticle formulations tested in vitro and in vivo. Intratracheal (IT) administration of these nanoparticles demonstrated that PBAE-PEG/lipids selectively targeted pulmonary epithelial cells, including alveolar type II cells. We compared RNA delivery with our PBAE-Lipid nanoparticles with PEI-based polyplex nanoparticles and commercial Lipofectamine, demonstrating highly active transfection efficiency in vitro and in vivo with the PBAE lipid nanoparticles.

In contrast, PBAE-LNP were more selective for delivery to airway and pulmonary epithelial cells through IT injection. Intravenous (IV) injection of the nanoparticles selectively delivered RNAs to pulmonary endothelial cells. We then utilized IV and IT delivery of Cre-recombinase (CRE-sRNA nanoparticles) to demonstrate cell type-specific transfection in B6.Cg-Gt(ROSA)26Sor^tm14(CAG-tdTomato)Hze^/J mice in vivo.

## Materials and Methods

### 2.1. Materials, DNA plasmids and mRNA

All lipids were purchased from Avanti Polar Lipids Lipids (Alabaster, AL, USA) including 18:1 TAP(DOTAP) (cat: 890890) and 18:1 DOPE (cat: 850725). 4A3-SC8(N-1438) was purchased from Echelon Biosciences (Salt Lake City, UT, USA). 1,4-butanediol diacrylate (cat: 411744), 4-amino-1-butanol (cat: 178330), citric acid (cat: 1.93427), 1,3-diaminopropane (cat: D23602), dexamethasone (cat: D4902) and ALT Activity Assay (cat: MAK052) were purchased from Sigma-Aldrich (St. Louis, MO, USA). D-Luciferin Bioluminescent Substrate (cat: 770504) was purchased from Revvity (Waltham, MA, USA). Luciferase-expressing plasmid DNA (pUL3; LOT 94344) was from Aldevron (Fargo, ND, USA). CleanCap® Cre mRNA (cat: L-7211) and CleanCap® Renilla Luc mRNA (5moU)(cat: L-7204) were from TriLink, San Diego, CA, USA).

### 2.2. PBAEs synthesis

Synthesis of the PBAE polymer backbone was conducted via a two-step Michael addition reaction, where 1,4-butanediol diacrylate and 4-amino-1-butanol were combined in a 1.1:2 molar ratio and reacted at 90°C for 24 hours to produce uncapped PBAE polymers[27]. These polymers were subsequently precipitated three times in cold ether to eliminate any unreacted materials, then dried under vacuum and lyophilized. For the capping of the terminal acrylate groups, the PBAE polymers were solubilized in tetrahydrofuran (THF) at a concentration of 100 mg/mL, and 30 molar equivalents of 1,3-diaminopropane were introduced. The reaction was carried out with stirring at room temperature for 4 to 5 hours. Finally, the end-capped PBAE polymers were isolated by precipitation in cold ether to remove any unreacted groups, then dried under vacuum and lyophilized[21]. Methoxy-PEG-succinimidyl succinates were then conjugated to the two terminal primary amine groups on either end of the 1,3-diaminopropane-capped PBAE polymer chain. This process involved transferring the synthesized PBAE polymers along with 2.05 molar equivalents of 5 kDa methoxy-PEG-succinimidyl succinate into a glass vial, which was then evacuated and filled with nitrogen. The reactants were dissolved in THF, and the mixture was stirred at room temperature overnight. The resultant PBAE-PEG polymer was then precipitated and washed three times with cold ether, followed by drying under vacuum and lyophilization. PBAE polymers were stored at -20 °C[21].

### 2.3. PBAE-lipid nanoparticle formulation

LNP formulations were prepared using the ethanol dilution method, where all lipids (4A3-SC8, DOPE, cholesterol and DOTAP) along with PBAE and PBAE-PEG were dissolved in ethanol at predetermined molar ratios. Concurrently, plasmid DNA or mRNA was dissolved in a 10 mM citrate buffer at pH 4.0. The lipids in ethanol and the plasmid DNA or RNA in the citrate buffer were then rapidly mixed using vortex at an ethanol-aqueous volume ratio of 1:3. This mixture was incubated at room temperature for 10 minutes and then dialyzed in PBS at pH 7 overnight, which achieves a final weight ratio of 30:1 (total lipids to mRNA or pDNA). The PEGylated LNP formulation is comprised of a lipid nanoparticle composition that includes 4A3-SC8, DOPE, cholesterol, PBAE-PEG, and DOTAP in a molar ratio of 20/20/35/2.5/60 respectively. The non-PEGylated LNP formulation consists of 4A3-SC8, DOPE, cholesterol, PBAE, and DOTAP, also in a molar ratio of 20/20/35/2.5/60. The only difference between the two formulations is the substitution of PBAE-PEG with PBAE in the non-PEGylated LNP composition[28].

### 2.4. Lipid-PBAE nanoparticles characterization

The hydrodynamic diameter and surface potential of the lipid-PBAE nanoparticles were measured using dynamic light scattering (DLS) on a Zetasizer (Malvern, Malvern, UK). Additionally, the structural properties of the nanoparticles were characterized by 1H NMR spectroscopy, performed in deuterated chloroform using a Bruker AV 400 MHz spectrometer (Billerica, MA, USA).

### 2.5. *In vitro* HEK293T cell transfections

The HEK293T cell line was cultured in Dulbecco’s Modified Eagle Medium (DMEM) (ThermoFisher, Waltham, MA, USA) enriched with 10% fetal calf serum (FCS) (ThermoFisher) and 1% Penicillin/Streptomycin (P/S) (ThermoFisher). Cell cultures were housed in a controlled environment at 37 °C and 5% CO2. Luciferase assays were performed as previously conducted[29] except that luciferase activity generated by PEGylated LNP or non-PEGylated LNP with 5-methoxyuridine modified or unmodified luciferase mRNA was compared with that generated by Lipofectamine RNAiMAX with the mRNA.

### 2.6. H441 ALI cultures and transfections

Air-liquid interface (ALI) cultures were initiated by seeding approximately 1 × 10^5^ cells per well in 12-well Transwell inserts (Corning), using base media for monocultures, and 5 × 10^4^ H441 cells in the case of co-cultures. The cells were allowed to attach and proliferate within the Transwells for 48 hours. On the third day, the apical medium was removed to facilitate air-lifting of the cells, simulating an air-liquid interface. Concurrently, the basolateral medium was replaced with either a base medium or a specialized polarization medium. This polarization medium, used for H441 cells and co-cultures, was composed of RPMI-1640 supplemented with 2 mM L-glutamine, 50 U/mL penicillin, 50 mg/mL streptomycin, 1% insulin-transferrin-selenium (GIBCO), 4% fetal calf serum (FCS), and 1 μM dexamethasone (Sigma). Media changes were conducted three times per week for the duration of the experiments. The cells were cultured under human surfactant air-liquid interface (SALI) conditions for 14 days post air-lift before any experimental procedures were conducted. The results are generated by EVOS fluorescence microscope.[30][31]

### 2.7. qRT-PCR

Quantitative real-time PCR (qRT-PCR) was employed to measure the RNA expression levels of surfactant protein C (SFTPC; Hs00951326_g1) and ATP-binding cassette sub-family A member 3 (ABCA3; Hs00184543_m1) in established air-liquid interface (ALI) cultures. RNA extraction from these cultures was performed using a QIAGEN RNeasy Mini Kit (QIAGEN). Following extraction, reverse transcription reactions were carried out with an iScript cDNA synthesis kit (Bio-Rad), adhering to the provided manufacturer’s instructions. The synthesized cDNA was then analyzed by quantitative PCR (qPCR) using a StepOnePlus Real-Time PCR System and specific TaqMan gene expression assays for STFPB and ABCA3 (Applied Biosystems). Expression levels of STFPB and ABCA3 were normalized against 18S RNA (TaqMan probe 4352930) to control for variations in RNA input and cDNA synthesis efficiency. The resultant data were expressed as mean ± standard error of the mean (SEM) and analyzed statistically using a 2-tailed Student’s t-test to determine significance. All statistical analyses were conducted using GraphPad Prism software, version 10 (GraphPad Software, Inc.).[32]

### 2.8. Animal studies

All animal experiments conducted were approved by the Institutional Animal Care and Use Committee (IACUC). B6.Cg-Gt(ROSA)26Sor^tm14(CAG-tdTomato)Hze^/J(Ai14) Cre reporter mice, FVBN mice and Nu/J mice aged 8–12 weeks were obtained from Jackson Laboratory. PEGylated LNPs with 5-methoxyuridine modified Cre mRNA (5moU) were administered to Ai14 tdTomato mice by either intratracheal (IT) administration or intravenous (IV) injection. Lungs were harvested five days after the LNP administration containing mRNA. Single cell suspensions were made from the lungs or lung sections were prepared. For *ex-vivo* imaging, FVBN mice were injected IV or IT with PEGylated LNPs with 5-methoxyuridine modified (5moU) or unmodified luciferase mRNA. Twenty-four hours after injection, mice received an intraperitoneal injection of 200 μL D-luciferin (30 mg/mL) and were euthanized 10 minutes later and internal organs such as heart, liver, spleen, lung, and kidney were collected for luminescence imaging. *In vivo* imaging studies utilized NU/J hairless mice injected with the LNPs with the mRNA (5moU) and D-luciferin as described above. To assess *in vivo* luciferase activity, we used IVIS Lumina III and Spectrum CT In Vivo Imaging Systems (PerkinElmer, Waltham, MA, USA). Whole-body imaging was performed 10 minutes after D-luciferin administration and luminescence intensity was quantified using Living Image software (Spectral Instruments, Tucson, AZ, USA).[22] Plasma alanine aminotransferase (ALT) activity, an indicator of liver function, was measured using a Diagnostic ALT test kit from Sigma-Aldrich (Catalog No. MAK052) according to the manufacturer’s instructions.[33]

### 2.9. Flowcytometry

Lung from Ai14 mice were digested in PBS medium with 480 U/ml Collagenase Type 1(Sigma Aldrich), 5U/ml Dispase(Sigma Aldrich) and 0.025 mg/ml DNase1(New England Biolabs, Ipswich, MA,USA). The red blood cells were removed using RBC lysis buffer(Biolegend, San Diego, CA, USA) for 5 min on ice. Single cell suspensions from lungs (2 × 10^6^ cells/well, 96 well plate) were stained with an antibody cocktail for 30 minutes on ice. The following antibodies were used for surface staining; CD45 (clone: I3/2.3, Biolegend, San Diego, CA, USA), CD31 (clone: 390, Invitrogen, Waltham, MA, USA), CD326 (clone: G8.8, Invitrogen) and CD140a (clone: APA5, Invitrogen). To exclude dead cells from the analysis, the LIVE/DEAD Fixable Near-IR Dead Cell Stain Kit (Invitrogen) was used. After surface staining, cells were washed 3 times with PBS containing 2% Fetal Bovine Serum (Corning, Glendale, AZ). Samples were acquired on a BD LSRFortessa (BD Biosciences, Franklin Lakes, NJ), and data were analyzed using FlowJo (BD Biosciences).[34]

### 2.10. Immunofluorescence

Lungs harvested from Ai14 tdTomato mice were cryosectioned at 5 μm for immunofluorescence studies, following a previously established protocol[5]. Specific cell types within these sections were identified using a series of primary antibodies: 1:500 rabbit anti-ERG (Roche, Basel, Switzerland) for endothelial cells, 1:1000 rabbit anti-NKX2.1 (Seven Hills, Cincinnati, OH,USA) for epithelial cells, 1:50 rabbit anti-MEOX2 (Atlas Antibodies, Bromma, Sweden) for fibroblasts, 1:500 rabbit anti-LAMP3 (OriGene, Rockville, MD, USA) for alveolar type 2 (AT2) cells, 1:500 mouse anti-SCGB1A1 (Seven Hills) for club cells, 1:500 rabbit anti-TUBA1A (Sigma Aldrich) for ciliated cells, and 1:200 goat anti-tdTomato (OriGene) to detect tdTomato expression. Sections were incubated with labeled secondary antibodies (Invitrogen) to visualize the primary antibody binding: donkey anti-rabbit Alexa Fluor (AF) 555 and AF 647 for rabbit primary antibodies, donkey anti-mouse AF 488 for mouse primary antibodies, and donkey anti-goat AF 555 for goat primary antibodies. Fluorescence images were captured using a Nikon A1R confocal microscope. The captured images were analyzed using Imaris software (company)

## 3. Results

### 3.1. Characterization of PBAEs synthesis and formulation of lipid-PBAE nanoparticles

We used two Poly(β-amino esters) (PBAEs) polymers, PBAE and PBAE-PEG to produce lipid-PBAE nanoparticles. PBAE polymers were synthesized through a step-growth polymerization process employing Michael addition to facilitate the reaction between 1,4-butanediol diacrylate and 4-amino-1-butanol. Hydrophilic 1,3-diaminopropane capping agents were applied to the hydrophobic PBAE backbones to create amphipathic polymers. (Fig. 1A). Then, PEGylated PBAE was made by using methoxy-PEG-succinimidyl succinates, which were reacted with the terminal primary amine groups at both ends of the 1,3-diaminopropane PBAE polymer chain (Fig. 1A)[21]. NMR demonstrated the completion of all reactions. The molecular peak at 1.7 and the peak at 4.2 are consistent with the PBAE backbone, and the k peak (Purple) with PBAE-PEG (Figure 2A). The amphipathic polymers PBAE and PBAE-PEG were then combined with other amphipathic lipids (4A3-SC8/DOPE/DOTAP/cholesterol) to form lipid-PBAE nanoparticles, encapsulating DNA or RNA to assemble the PBAE/LNP.(Fig. 1B, C). We assessed the morphological characteristics of PBAE-PEG/LNP and PBAE/LNP with transmission electron microscopy (TEM). Lipid-PBAE nanoparticles were typical of a mesoporous structure, distinct from the dual-layered structures typical of standard lipid nanoparticles(4A3-SC8/DOPE/DOTAP/ DEM-PEG/cholesterol)(Figure 2B). To predict how PEGylated PBAE (PBAE-PEG) lipid nanoparticles release DNA or mRNA after entering into cells, we measured the size of PBAE-PEG/LNP in PBS at pH 7 and 10 μM or 100 μM citric acid buffer at pH 6 and pH 5. At a neutral pH 7, the nanoparticles exhibited an average size of 108.1 nm (PDI=0.140). As the pH decreased, the size increased progressively from 148.4 nm (PDI=0.154) at pH 6 to 294 nm (PDI=1.000) at pH 5. The size distribution indicated that PEGylated PBAE (PBAE-PEG) lipid nanoparticles were unstable at pH 5 since there were two significant peaks at 20.16 nm (28.2%) and 294 nm (71.8%) (Fig. 2C), suggesting that PEGylated PBAE (PBAE-PEG) lipid nanoparticles may release DNA or mRNA in the endocytosis process at low pH.

**Fig. 1.**
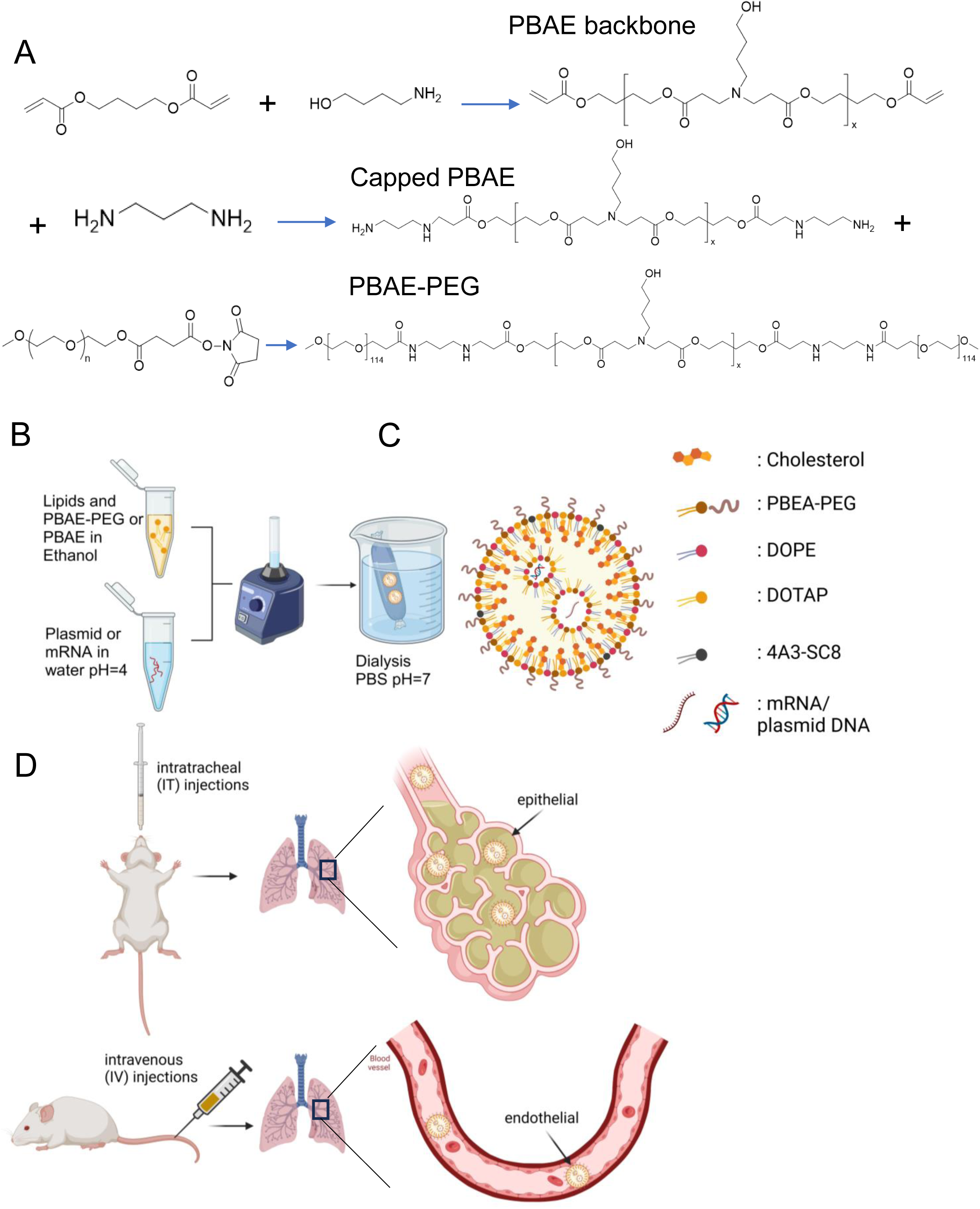
Overview of the PBAE synthesis, PBAE/LNP formulation and cell specific gene delivery to the lung. (A) Uncapped PBAE polymers were synthesized by the Michael addition reaction of 1,4-butanediol diacrylate and 4-amino-1-butanol to produce the PBAE backbone. Subsequently, uncapped PBAE polymers were capped with 1,3-diaminopropane (Capped PBAE). PBAE-PEG (PEGylated PBAE) was synthesized by conjugating methoxy-PEG-succinimidyl succinates to capped-PBAE at a 2:1 PEG-to-PBAE molar ratio. (B) Plasmid DNA (or mRNA) dissolved in water at pH 4 was assembled by vortex mixing in non-PEGylated PBAE (PBAE) or PEGylated PBAE (PBAE-PEG)/4A3-SC8/DOPE/Cholesterol/DOTAP lipid nanoparticles (LNPs) dissolved in ethanol and then dialyzed in PBS at pH 7 overnight. The figure drawing was created by BioRender.com. (C) Shown is a schematic of PEGylated PBAE (PBAE-PEG)/4A3-SC8/DOPE/Cholesterol/DOTAP lipid nanoparticles (LNPs) carrying plasmid DNA (or mRNA). The figure drawing was created by BioRender.com. (D) Shown are schemes demonstrating different routes of injections (intravenous [IV; top panel] vs. intratracheal [IT; bottom panel]) with LNPs with mRNA. The figure drawing was created by BioRender.com.

**Fig. 2.**
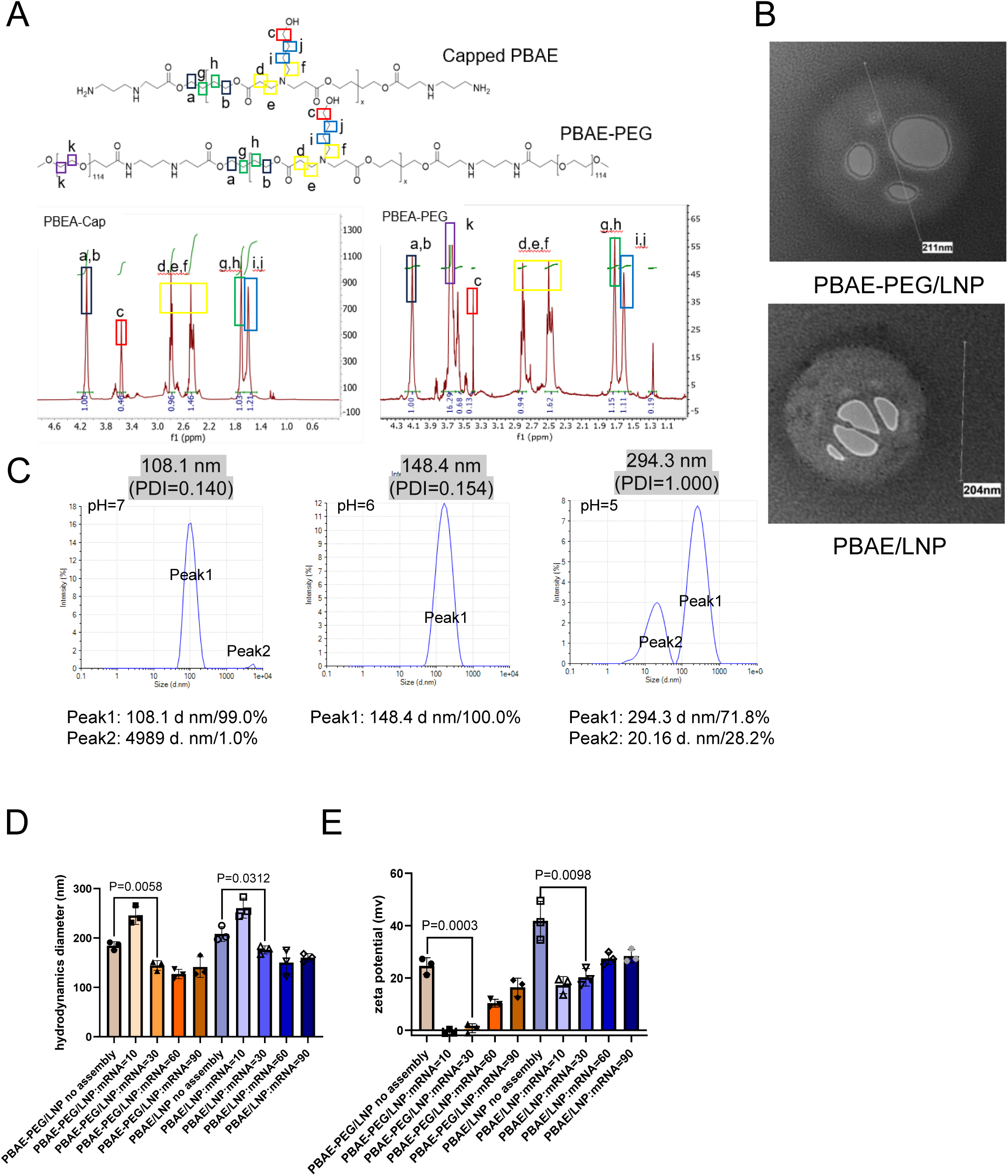
PBAE/LNP and PBAE-PEG/LNP characterization. (A) Shown are chemical structures (top panel) and 1H NMR spectra (bottom panel) of Capped PBAE (Capped PBAE) and PBAE-PEG (PEGylated PBAE). PEGylation is demonstrated by peak k (purple). (B) Shown are representative transmission electron microscopy (TEM) images of PBAE-PEG/4A3-SC8/DOPE/Cholesterol/DOTAP lipid nanoparticles (LNPs) carrying plasmid DNA (top panel) and PBAE/4A3-SC8/DOPE/Cholesterol/DOTAP lipid nanoparticles (LNPs) carrying luciferase-expressing plasmid DNA (bottom panel) with the nanoparticle diameter of 211 nm and 204 nm, respectively (mean value from n=3 fields of view). (C) Shown are the diameter of PEGylated PBAE/4A3-SC8/DOPE/Cholesterol/DOTAP lipid nanoparticles (LNPs) carrying plasmid DNA assembled in PBS at pH 7 (left panel), citric acid buffer at pH 6 (middle panel) and pH 5 (right panel). (D) Shown is the size (hydrodynamics diameter) of PEGylated (PBAE-PEG) and non-PEGylated (PBAE) LNPs assembled with mRNA at different mass ratio (LNPs:mRNA = 10:1 to 90:1). (E) Shown is the zeta potential of PEGylated (PBAE-PEG) and non-PEGylated (PBAE) LNPs assembled with mRNA at different mass ratios (LNPs:mRNA = 10:1 to 90:1).

### 3.2. In vitro transfection validation PBAE-PEG/LNP, PBAE/LNP, PEI-based nanoparticles, and Lipofectamine

In order to determine the best mass ratio between PEGylated or non-PEGylated PBAE/4A3-SC8 lipid nanoparticles (hereafter, LNP) and mRNA for transfection efficiency, we tested different mass ratios from 10:1 to 90:1 (LNP:mRNA) using luciferase mRNA in HEK293T cells *in vitro*. Ratios of 30:1 and 60:1 provided the highest transfection efficiency as measured by luciferase activity compared to those of 10:1 or 90:1 (Fig. 3F). The size and surface charge of PEGylated (PBAE-PEG) or non-PEGylated (PBAE) LNP with different amounts of the mRNA was also determined by Zetasizer. The size of the PEGylated (PBAE-PEG) LNP itself was 184.3 nm. When assembled with the mRNA at the mass ratio of 10:1 (LNP:mRNA), the size significantly increased to 245.6 nm. When assembled with mRNA at the mass ratio of 30:1 or higher (LNP:mRNA), the size significantly decreased to below 144.3 nm (Fig. 2E). On the other hand, the size of non-PEGylated (PBAE) LNP itself was 208 nm. When assembled with the mRNA at the mass ratio of 10:1 (LNP:mRNA), the size significantly increased to 260.3 nm. When assembled with the mRNA at the mass ratio of 30:1 or higher (LNP:mRNA), the size significantly decreased to below 176 nm(Fig. 2D). The zeta potential of PEGylated (PBAE-PEG) or non-PEGylated (PBAE) LNP itself was 24.3 mV or 41.3 mV, respectively. When assembled with the mRNA at the mass ratio of 30:1, the zeta potential of PEGylated (PBAE-PEG) or non-PEGylated (PBAE) LNP was 2.2 mV and 27 mV, respectively. When assembled with the mRNA at the mass ratio of 60:1 or 90:1, the zeta potential of PEGylated (PBAE-PEG) or non-PEGylated (PBAE) LNP was higher than 5.67 mV and 1 mV, respectively(Fig. 2E). Considering that the recommended size and charge of lipid nanoparticles are < 200 nm and ∼20 mV for targeting lung cells[35], [36]and using our luciferase data (Fig. 3F), we chose the mass ratio of 30:1 (LNP:nucleic acid) for the further experiments below. We tested the efficiency of transfection of PBAE-PEG/LNP and PBAE/LNP formulations and compared these with PEI-PEG-LinA previously described[37] and Lipofectamine 3000 in HEK 293T cells the transfection activity was similar after first 24 hours; but PBAE-PEG/LNP been detect the most of tdTomato fluorescence after 48 hours(Figure 3A). In preparation for in vivo studies, we evaluated the efficacy of PBAE-PEG/LNP and PBAE/LNP for mRNA delivery, quantifying luciferase activity, RNAiMAX serving as a positive control. PBAE-PEG/LNP and PBAE/LNP were significantly more active than RNAiMAX (Fig. 3D). We tested PBAE-PEG/LNP and PBAE/LNP in an air-liquid interface (ALI) culture of H441 cells (a human non-small cell adenocarcinoma cell line)[30] q-PCR shows after airlifting that STFPB and ABCA3 expression increased(Fig. 3F). Compared with Lipofectamine 3000, PBAE-PEG/LNP was more active (Fig. 3B).

**Fig. 3.**
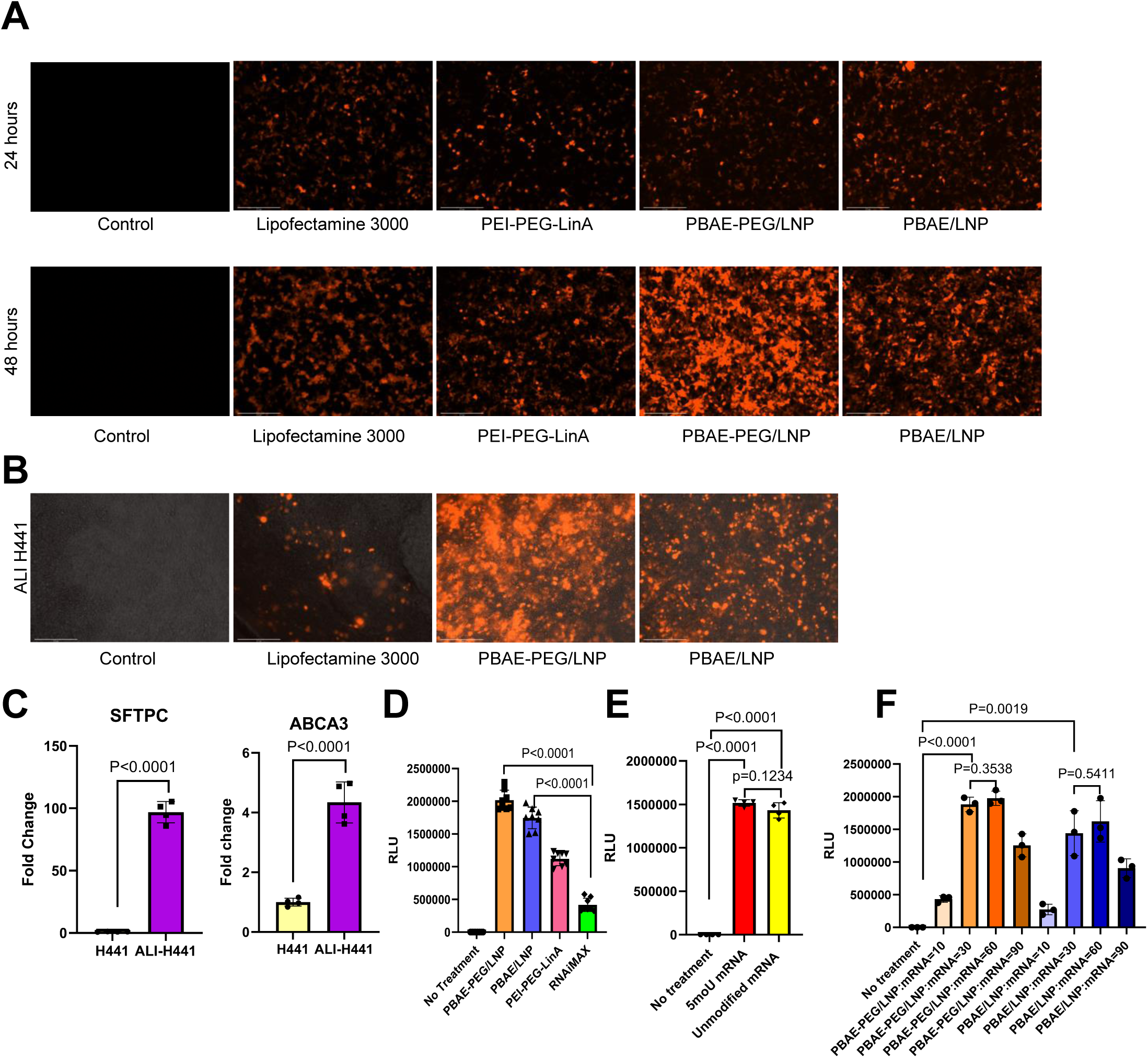
In vitro activity of LNPs and PEI-PEG-LinA nanoparticles in HEK293T and H441 cells at air-liquid-interface(ALI) (A) EVOS fluorescence microscopy shows the activity of gWiz-mcherry plasmid with PBAE-PEG/LNP, PBAE/LNP, and PEG-LinA nanoparticles on HEK293T cells, 24 and 48 hours after transfection. Scale bar=275μm. (B) EVOS fluorescence microscopy shows the activity of gWiz-mcherry plasmid with DOPE/DSPE-PEG/DOTAP/Chol, PBAE-PEG/LNP, and PBAE/LNP on H441 air-liquid interface (ALI) 4 days after transfection. (C) *ABCA3* and *SFTPC* mRNA in normal H441 cells and ALI H441 cells 11 days after airlifting. normalized to β-actin. Results are expressed as mean SD of replicates for each group (n=4, unpaired, two-tailed Student’s t-test). (D) Shown is luciferase activity (RLU) from HEK293T cells transfected with PBAE-PEG/LNP, PBAE/LNPs, PEI-PEG-LinA or Lipofectamine (RNAiMAX) carrying unmodified luciferase mRNA (2 μg) at the mass ratio of 30:1 (LNPs:mRNA) for LNPs as instructed by the manufacturer for RNAiMAX. (E) Shown is luciferase activity (RLU) from HEK293T cells transfected with PEGylated (PBAE-PEG) carrying 5moU or unmodified luciferase mRNA (2 ug) at the mass ratio of 30:1 (LNPs:mRNA). (F) Shown is luciferase activity (RLU) from HEK293T cells transfected with PEGylated (PBAE-PEG) and non-PEGylated (PBAE) LNPs carrying unmodified luciferase mRNA (2 μg) at different mass ratios (LNPs:mRNA = 10:1 to 90:1). Results are expressed as mean SD of replicates for each group (n=3, unpaired, two-tailed Student’s t-test).

### 3.3. Biodistribution analysis of PBAE-PEG/LNP, PBAE/LNP after intravenous and intratracheal administration

We compared 5moU mRNA and unmodified mRNA for gene delivery.[38] Measuring luciferase expression after transfected with 5moU mRNA and unmodified mRNA. Both 5moU mRNA and unmodified mRNA with PBAE-PEG/LNP showed similar Relative Light Unit (RLU) in HEK293T cell at 48 post-transfection (Figure 3E). We compared PBAE-PEG/LNP luciferase 5moU mRNA versus unmodified mRNA in vivo delivery in FVB/N mice and found out 5moU mRNA has higher luciferase activity (Figure 4A, B), indicating that 5-methoxyuridine modification of mRNA is required for *in vivo* transfection regardless of the injection route (IV or IT). To study the in vivo biodistribution of gene transfer with PBAE-PEG/LNP and PBAE/LNP, we encapsulated luciferase 5moU mRNA and unmodified mRNA and tested these in Nude/J mice via IT and IV injections. Bioluminescence was assessed after dissecting various organs. Injection of PBAE-PEG/LNP with 5moU luciferase mRNA via IT injection resulted in luciferase expression in the lung only. After IV injection, nanoparticle luciferase activity was detected in the lung, liver, and spleen(Figure 4C).

**Fig. 4.**
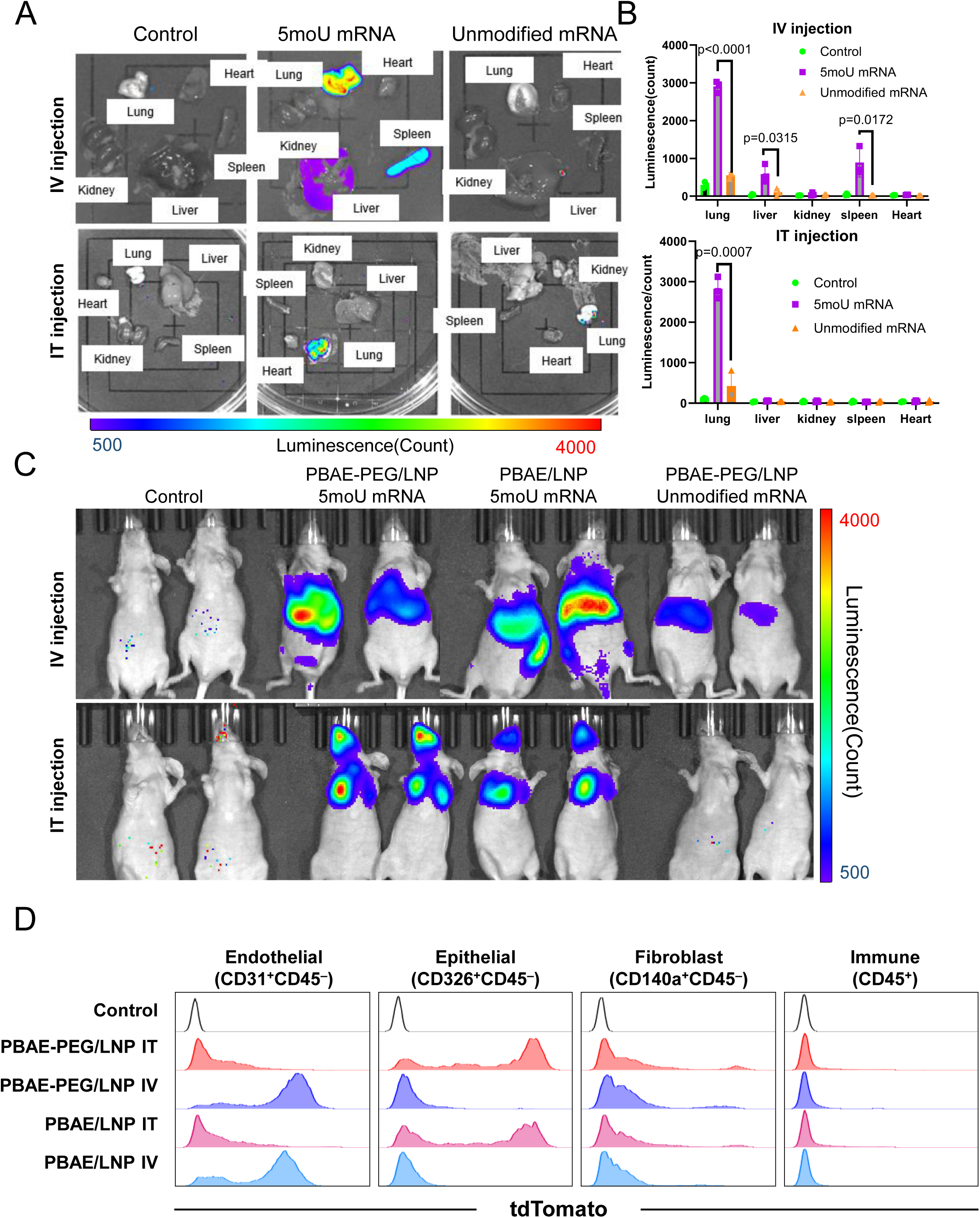
Activity of PBAE-PEG/Lipids and PBAE/Lipids nanoparticles for delivery of mRNA and stabilized mRNA in vivo. (A) Shown are representative luminescent ex vivo images at post-transfection 24 hours detecting the biodistribution of luciferase activity from different organs (lung, spleen, liver, heart and kidney) of FVB/N mice (n=3/group) after IV (top panel) or IT (bottom panel) injections of PBAE-PEG/LNP with 10μg/mice of 5moU or unmodified luciferase mRNA (B) Shown are quantification of luminesce images by ImageJ from 3 mice/group as described in (A) Results are expressed as mean SD of triplicates for each group (unpaired, two-tailed Student’s *t*-test). (C) Shown are representative luminesce images detecting the biodistribution of luciferase activity as described in (A) except that whole bodies of nude mice (n=4) were visualized by IVIS. (D) FACS histograms show tdTomato expression in endothelial cells (CD31^+^CD45^−^), epithelial cells (CD326^+^CD45^−^), lung fibroblasts (CD140a^+^CD45^−^), and immune cells (CD45^+^) from lungs of tdTomato mice transfected with PBAE-PEG/LNP and PBAE/LNP carrying 5-methoxyuridine modified Cre mRNA (5moU). Results were representative from 2 independent experiments with 2 or 3 mice per group.

### 3.4. Pulmonary cell type selectively activity of PBAE-PEG/LNP, PBAE/LNP after IT instillation

To analyze LNPs in vivo transfection after IT injection, we use FACS to analyze the Ai14 mice treated by PBAE-PEG/LNP and PBAE/LNP with 5moU Cre-recombinase mRNA. The expression of Cre-recombinase causes the deletion of a stop cassette that induces the selective cell expression of td-Tomato in transfected cells. Flow cytometry data demonstrated that 69.25% and 45.6% of pulmonary epithelial cells were transfected by PBAE-PEG/LNP and PBAE/LNP, respectively (Supplemental Fig. 2A, B). IT injection of PBAE-PEG/LNP and PBAE/LNP would only be transfected lung epithelial cells (CD326+CD45–)(Fig. 4D). We utilized nanoparticle preparations of PBAE-PEG/LNP and PBAE/LNP to deliver 5moU Cre-recombinase mRNA to identify cell type-specific expression sites after IT administration in vivo. Targeting was identified by fluorescence microscopy to identify td-Tomato co-expression with cell-type selective pulmonary cells. PBAE-PEG/LNP and PBAE/LNP selectively expressed Cre-recombinase mRNA in pulmonary epithelial cells identified by co-staining td-Tomato and NKX2.1, a cell-specific marker for pulmonary epithelial cells (Fig. 5A). td-Tomato was co-expressed with LAMP3, a known AT2 cell-selective marker (Fig. 5A)[39]. PBAE-PEG/LNP was active in both alveolar and bronchial epithelial cells, while PBAE/LNP was more active in conducting airway epithelial cells. Co-staining of td-Tomato with MEOX2, a pulmonary fibroblast cell maker, was not detected after IT injection (Fig. 5C). Likewise, co-staining of td-Tomato with ERG, a pulmonary endothelial cell marker was not observed (Fig. 5B)[40]. To identify airway epithelial cell types targeted by PBAE-PEG/LNP and PBAE/LNP, we co-stained td-Tomato with SCGB1A1 a club cell marker [41] and TUBA1A a pulmonary ciliated cells marker[42]. Recombination was observed in club cells marked by Scgb1a1 but not in ciliated cells marked by TUBA1A (Figure 5D).

**Fig. 5.**
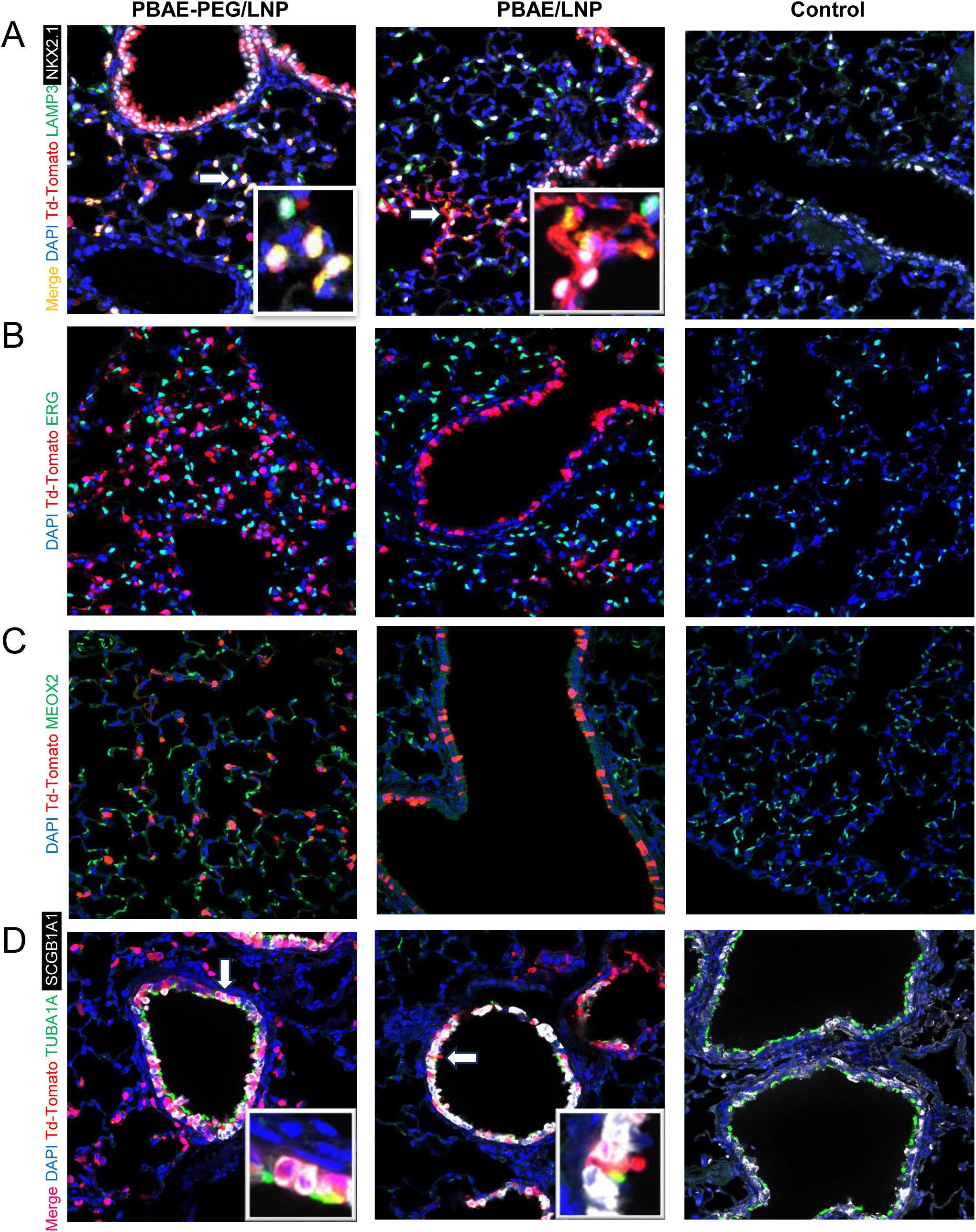
Confocal immunofluorescence microscopy of lung sections shows cell type specification by IT administration of PBAE-PEG/Lipids and PBAE/Lipids nanoparticles. Cre-recombinase stabilized mRNA encapsulated by PBAE-PEG/LNP and PBAE/LNP were administrated to B6.Cg-Gt(ROSA)26Sortm14(CAG-tdTomato)Hze/J (Ai14) mouse by IT injection after 4 days treatment. Lung tissue was harvested and sectioned to prepare for immunofluorescence staining for confocal microscopy to identify cell selectivity. n=3. (A) Td-Tomato (red) fluorescence was co-expressed with pulmonary epithelial cell marker NKX2.1 (white) and alveolar epithelial Type II(AT2) cell marker LAMP3 (green). Cell nuclei were counterstained with DAPI (blue). (B) Td-Tomato (red) fluorescence was not co-expressed with endothelial cell marker ERG (green). Cell nuclei were counterstained with DAPI (blue). (C) Td-Tomato (red) fluorescence was not co-expressed with fibroblast marker MEOX2 (green). Cell nuclei were counterstained with DAPI (blue). (D) Td-Tomato (red) fluorescence was co-expressed with club cell marker SCGB1A1 (white). td-Tomato (red) fluorescence not co-expressed with ciliated-cell marker TUBA1A (green). Cell nuclei were counterstained with DAPI (blue).

### 3.5. Pulmonary cell type selective activity of PBAE-PEG/LNP, PBAE/LNP after IV instillation

To identify our LNPs in vivo transfection after IV injection we use FACS to analyze the Ai14 mice treated by PBAE-PEG/LNP and PBAE/LNP, 5moU Cre-recombinase mRNA. Flow cytometry data demonstrated that approximately 76.7% and 67.1% of pulmonary endothelial cells were transfected PBAE-PEG/LNP and PBAE/LNP, respectively (Supplemental Fig. 2A, B). The FACS study shows IV injection of PBAE-PEG/LNP and PBAE/LNP would only be transfected lung endothelial cells (CD31+CD45–)(Fig. 4D). We utilized nanoparticle preparations of PBAE-PEG/LNP and PBAE/LNP to deliver stabilized Cre mRNA to activate td-Tomato expression after intravenous administration to adult mice. td-Tomato was co-expressed with ERG, a cell-selective marker for pulmonary endothelial cells after IV administration (Fig. 6B). Co-staining of td-Tomato with MEOX2, a pulmonary fibroblast cell maker, was not detected (Fig. 6C). Likewise, td-Tomato was not co-expressed with NKX2.1 or LAMP3 (Fig. 6A). Co-expression of td-Tomato with SCGB1A1 or TUBA1A was not detected (Fig. 6D).

**Figure 6.**
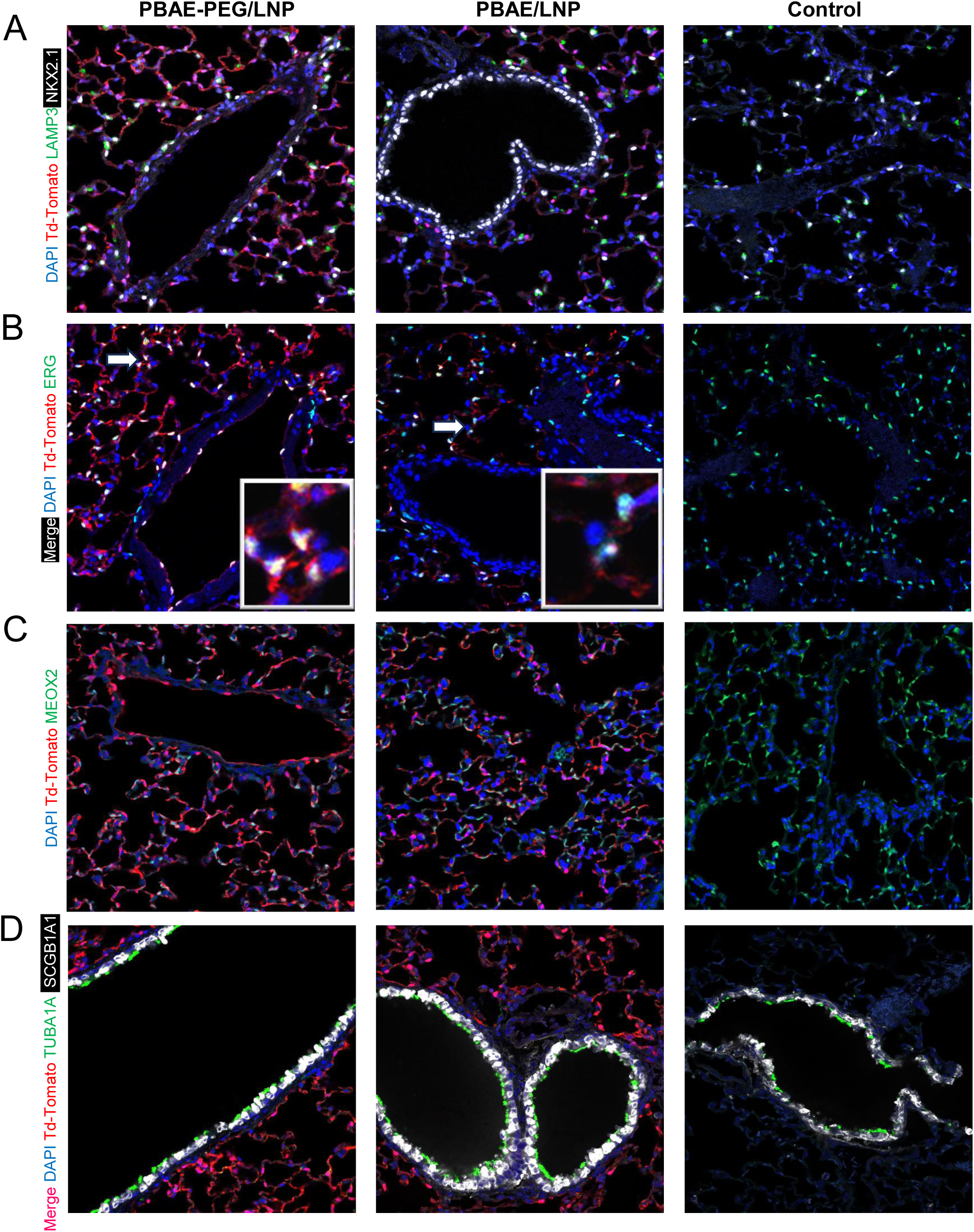
Confocal immunofluorescence microscopy of lung sections shows cell type specification by IV administration of PBAE-PEG/LNP and PBAE/LNP nanoparticles. Cre-recombinase stabilized mRNA encapsulated by PBAE-PEG/LNP and PBAE/LNP to delivery to B6.Cg-*Gt(ROSA)26Sor^tm14(CAG-tdTomato)Hze^*/J (Ai14) mouse by IT injection for 4 days. Lung tissue was harvested and sectioned to prepare for immunofluorescence staining for confocal microscopy to identify cell selectivity. n=3. (A) Td-Tomato (red) fluorescence was not co-expressed with pulmonary epithelial cell marker NKX2.1 (white) and alveolar epithelial Type II(AT2) cell marker LAMP3 (green). Cell nuclei were counterstained with DAPI (blue). (B) Td-Tomato (red) fluorescence was co-expressed with endothelial cell marker ERG (green). Cell nuclei were counterstained with DAPI (blue) (C) Td-Tomato (red) fluorescence was not co-expressed with fibroblast marker MEOX2 (green). Cell nuclei were counterstained with DAPI (blue) (D) Td-Tomato (red) fluorescence was not co-expressed with club cell marker SCGB1A1 (white). td-Tomato (red) fluorescence not co-expressed with ciliated-cell marker TUBA1A (green). Cell nuclei were counterstained with DAPI (blue)

### 3.6. Toxicity study of LNPs in both In Vitro and In Vivo

To evaluate potential toxicity associated with PBAE-PEG/LNP and PBAE/LNP, we assessed alanine aminotransferase activity (ALT), a marker of liver toxicity after IT and IV administration of PBAE-PEG/LNP and PBAE/LNP, observing no changes in ALT (Supplemental Fig. 1A). PBAE-PEG/LNP and PBAE/LNP did not cause injury using in vitro MTS Assay, HEK293T cells (Figure 6B). Pulmonary histology was not altered after IT and IV injection of PBAE-PEG/LNP and PBAE/LNP 2 days after administration (Supplemental Fig. 1A).

## Discussion

While lipid nanoparticles COVID-19 vaccines were effective in reducing the spread of SARS-CoV-2, these vaccines do not require cell type-specific, sustained, or high levels of gene expression. In contrast, gene therapy faces significant challenges due to the limitations of gene delivery systems in achieving sustained gene expression without causing inflammation. Various viral vectors, including lentiviruses[43], and adeno-associated viruses (AAVs)[44], have been developed for gene transfer to pulmonary cells. However, targeting specific cell types, handling the size and complexity of therapeutic genetic cargoes, and mitigating inflammation and immune responses remain major obstacles. Lentiviruses, for instance, elicit immune responses that hinder lung gene therapy. AAVs cannot be administered repeatedly and cannot deliver large RNA or DNA sequences, making correction of pulmonary gene deficiencies a persistent challenge in gene therapy.

Recent advancements in nanoparticle development have shown significant promise. Technologies such as SORT lipid nanoparticles[26], [45], PFV-LNPs[46], and polymer-lipid nanoparticles[47], [48] present substantial potential for gene therapy. SORT LNPs, for instance, exhibit remarkable lung targeting ability, enhanced by increased DOTAP to improve lung targeting via IV administration and include helper lipids to enhance transfection efficiency. SORT LNPs have demonstrated both lung endothelial and epithelial transfection ability. Similarly, PFV-LNPs have shown effective delivery of mRNA to the retina, achieving up to 50% efficiency compared to regular LNPs. Polymer-lipid nanoparticles, which incorporate PBAE to control physiochemical characteristics like lipophilicity, show transfection efficacy several times higher than polyethylenimine (PEI).

Despite these advances, pulmonary cell-specific delivery systems remain challenging. In our study, we developed two novel nanoparticles, PBAE-PEG/LNP and PBAE/LNP, to deliver RNA into lung epithelial cells, including AT2 cells and airway club cells, via IT injection, and lung endothelial cells via IV injection. Our results demonstrated that PEGylation affects targeting ability. Specifically, PBAE-PEG/LNP with PEGylation delivered mRNA to lung epithelial cells, including AT2 cells, while PBAE/LNP without PEGylation targeted airway club cells via IT injection. PEGylation enhanced transfection efficiency via IV injection. Stabilized mRNA showed higher expression than naked mRNA for in vivo transfection, without causing liver toxicity and lung inflammation.

Surface charge is a critical parameter in the formulation of DNA or mRNA nanoparticles, as electrostatic interactions typically occur between cationic polymers and negatively charged nucleic acids. Studies suggest that lung-targeting nanoparticles should have a zeta potential of +20 to +50 mV[37], [49]. However, positive charges in cationic polymers often lead to cytotoxicity[50][51]. Our PBAE-PEG/LNP formulation demonstrated low zeta potential while effectively delivering genetic cargo via both IT and IV administrations. Typically, high surface charges can disrupt mitochondrial function and cell metabolism, posing significant toxicity risks[52][53]. PEGylation enhanced gene delivery to the lung during IV administration, reducing apolipoprotein E (ApoE) interaction with lipids, and facilitating low-density-lipoprotein (LDL) receptor-mediated cellular uptake by hepatocytes, thereby increasing delivery to the pulmonary vasculature [54], [55], [56]. Our study aligns with these findings, showing that PBAE-PEG/LNP has higher lung transfection efficiency than PBAE/LNP during IV injection. However, IV injection still led to high expression in the liver, perhaps due to interactions between ApoE and lipids. Our use of 4A3-SC8 [26], [45] as a helper lipid and a formulation similar to SORT nanoparticles for PBAE-PEG/LNP and PBAE/LNP, but incorporating PBAEs, enhanced cell-specific targeting abilities. SORT lipid nanoparticles demonstrated lung cell targeting capabilities, transfecting both lung epithelial cells and endothelial cells via IV injection[45]. Our work showed that IV injection selectively targeted lung endothelial cells and failed to cross the endothelial barrier to stromal or epithelial cells. IT injection of PBAE-PEG/LNP and PBAE/LNP had somewhat different targeting abilities, with PBAE-PEG/LNP delivering to epithelial cells, including AT2 cells, and PBAE/LNP targeting airway secretory (club) cells.

Despite reduced stability and increased clearance by alveolar macrophages, PBAE/LNP, which lacks PEGylation, selectively targeted airway club cells during IT administration. PBAE/LNP presents a promising alternative for specific pulmonary conditions where targeted gene delivery to airway club cells, which serve as progenitors of secretory and ciliated cells is needed. PBAE-PEG/LNP is considered a potential candidate for delivering therapeutic cargo to AT2 cells for genetic disorders, for example, ABCA3 deficiency. We utilized smRNA with uridines substituted with 5-methoxyuridine to reduce inflammation, consistent with findings that methylated mRNA reduces the activation of the toll-like receptor signaling [57], minimizing inflammatory responses post-administration via both IV and IT routes. In our present studies, we observed gene expression without cell toxicity or overt inflammatory responses after IV or IT delivery.

## 5. Conclusions

In summary, our work proposes that PBAE-lipid nanoparticles have different targeting abilities after IT or IV injection. We confirmed previous findings that smRNA is more effective than RNA for in-vivo gene delivery. Needed are further studies to assess the duration of expression and whether levels of protein expression will be an effective treatment of genetic disorders. Our study demonstrated selective gene transfer to pulmonary endothelial and epithelial cells, providing a framework for designing more permanent strategies for pulmonary gene therapy. With the successful IV injection delivery of LNP with sgRNA to mouse liver for gene editing[58], nanoparticle gene therapy for disorders caused by mutations in ABCA3, CFTR, SFTPB, and SFTPC, and other lung diseases caused by mutations is becoming increasingly promising.

## Acknowledgments

This work was supported by NIH grants (U01HL134745, U01HL148856, R01HL164414, R01CA240317, R01AI139675), University of Cincinnati Cancer Center Pilot Project Award Program grant (S323349-2023 Ride Cincinnati-Maeda), Cincinnati Children’s Hospital Medical Center (Trustee Award Grant, CF-RDP Pilot & Feasibility Grant, GAP funding). YS was supported by NIH T32 Vaccinology Training Program T32AI165396 and the Japan Society for the Promotion of Science Overseas Research Fellowships. We thank Iris Fink-Baldauf, Jaymi St. Arnold and Mary Durbin for assistance and discussions.

## CRediT authorship contribution statement

**Bingxin Liu:** Conceptualization, Methodology, Formal analysis, Investigation, Data curation, Writing. **Yamato Sajiki:** Methodology Data curation. **Anusha Sridharan:** Methodology. **William D. Stuart:** Methodology, Writing. **Xuemei Cui:** Methodology. **Matthew Siefert:** Methodology. **Koichi Araki:** Data curation, Funding acquisition. **Assem G. Ziady**: Data curation. **Donglu Shi:** Conceptualization, writing. **Jeffery A. Whitsett:** Conceptualization, Data curation, writing, Funding acquisition Investigation, Resources, Project administration. **Yutaka Maeda:** Conceptualization, Methodology, Data curation, writing, Funding acquisition, Investigation, Resources, Project administration, Validation, Formal analysis, Supervision.

---

**Supplemental Fig. 1.**
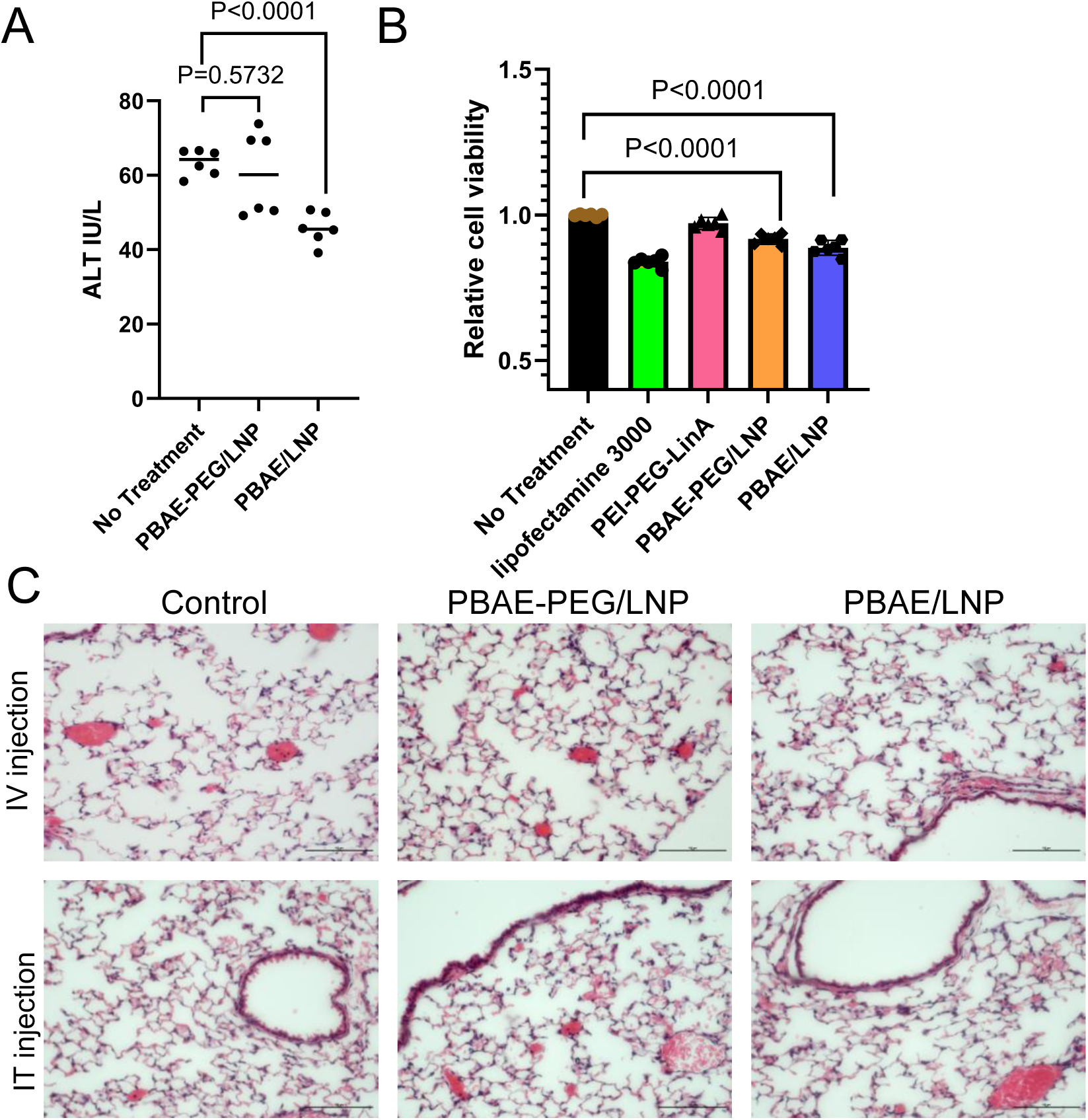
Toxicity study of PBAE-PEG/LNP and PBAE/LNP. (A) Alanine Aminotransferase Activity Assay (ALT) assay to determine toxicity of PBAE-PEG/LNP and PBAE/LNP for in vivo delivery. There is no significant liver toxicity after IV injections of 25ug mRNA LNP1 and LNP2 nanoparticles per mouse at post-transfection 4 days (n=4, unpaired, two-tailed Student’s t-test) (B) MTS assay indicates that PBAE-PEG/LNP and PBAE/LNP does not reduce cell viability in HEK293T cells. (n=6, unpaired, two-tailed Student’s t-test) (C) Lung histology in treated and untreated mice. Representative H&E stained lung sections (20×) at post-transfection 4 days of PBAE-PEG/LNP and PBAE/LNP. N=4/group

**Supplemental Fig. 2.**
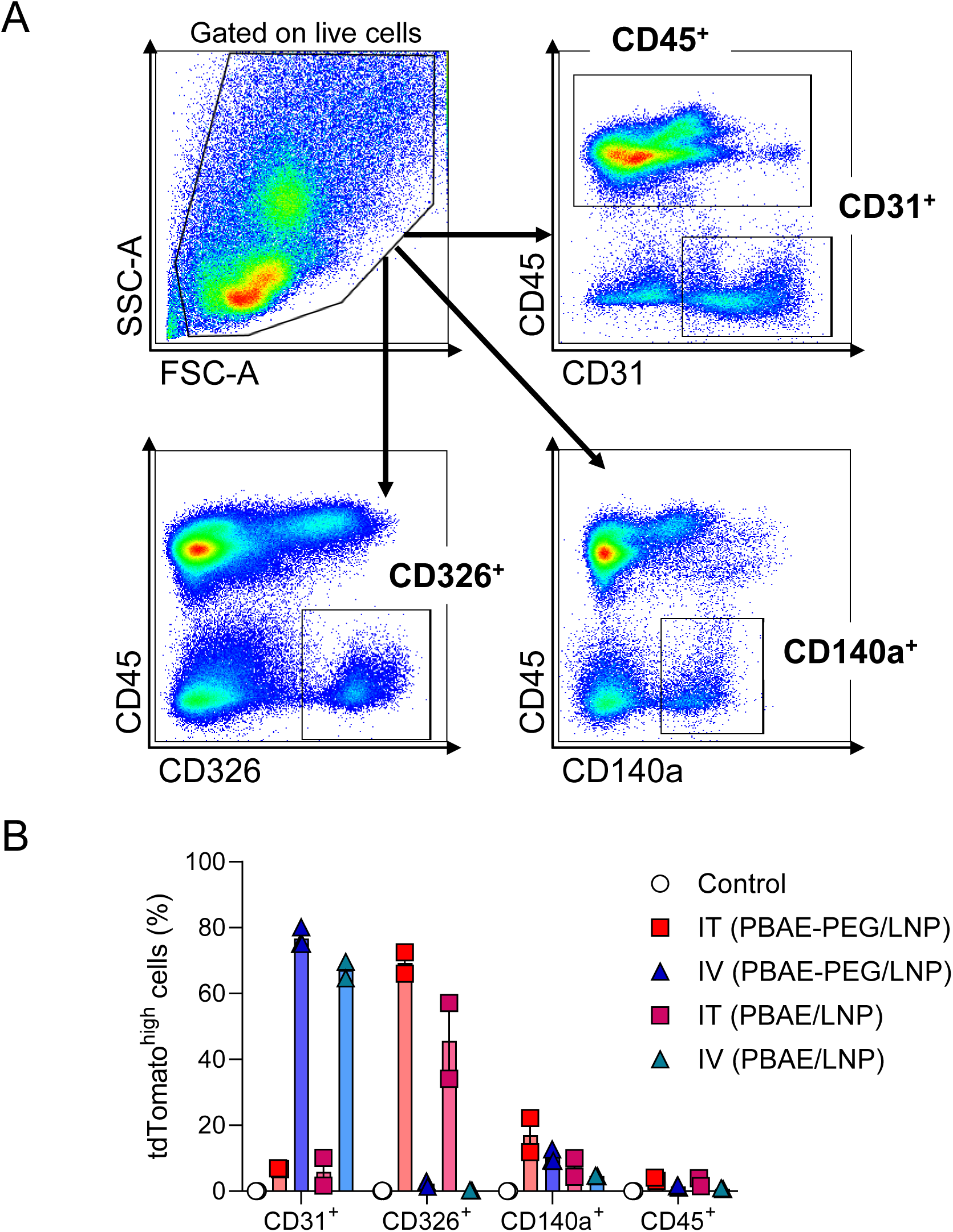
Gating strategy and quantified data of flowcytometry analysis. (A) Gating strategy for the identification of cell lineage populations (CD31^+^CD45^−^ as endothelial cells, CD326^+^CD45^−^ as epithelial cells, CD140a^+^CD45^−^ as fibroblasts, and CD45^+^ as immune cells) from whole-lung single cell suspension. (B) Graph displays the frequency of tdTomato^high^ cells in endothelial cells, epithelial cells, fibroblasts, and immune cells. Results were pooled from 2 independent experiments with 1 or 2 mice per group. Each symbol represents an individual mouse.

## Notes

### Competing Interest Statement

The authors have declared no competing interest.

